# Co-transcriptional loading of RNA export factors shapes the human transcriptome

**DOI:** 10.1101/318709

**Authors:** Nicolas Viphakone, Ian M Sudbery, Catherine G. Heath, David Sims, Stuart A. Wilson

## Abstract

During gene expression, RNA export factors are mainly known for driving nucleocytoplasmic transport. Whilst early studies suggested that the <u>E</u>xon <u>J</u>unction <u>C</u>omplex provides a binding platform for them, subsequent work proposed that they are only recruited by the <u>C</u>ap-<u>B</u>inding <u>C</u>omplex to the 5’ end of RNAs, as part of TREX. Using iCLIP, we show that the export receptor Nxf1 and two TREX subunits, Alyref and Chtop, are actually recruited to the whole mRNA co-transcriptionally via splicing but before 3’-end processing. Consequently, Alyref alters splicing decisions and Chtop regulates alternative polyadenylation. Alyref is recruited to the 5’-end of RNAs by CBC and our data reveal subsequent binding to RNAs near EJCs. We demonstrate eIF4A3 stimulates Alyref deposition not only on spliced RNAs close to EJC sites, but also single exon transcripts. Our study reveals mechanistic insights into the co-transcriptional recruitment of mRNA export factors and how this shapes the human transcriptome.

## INTRODUCTION

RNA polymerase II (RNAPII) transcribes most human genes as RNA precursors that mature through several stages to reach their functional state. These include 5’ capping, splicing, and 3’-end processing which involves 3’ endonucleolytic <u>c</u>leavage followed by <u>p</u>oly<u>a</u>denylation (CPA). This maturation also deposits RNA-binding proteins (RBPs) as completion marks for downstream events such as RNA nucleocytoplasmic transport, localization, translation, and stability (Singh et al., 2015).

Splicing is mostly co-transcriptional in humans (Tilgner et al., 2012), with ~65% introns removed from newly-made RNAs in only 5 minutes (Windhager et al., 2012). Yet, up to 80% of pre-mRNAs can be affected by at least one inefficient splicing event (Middleton et al., 2017), causing the retention of 5-15% of all expressed introns within mRNAs (Boutz et al., 2015). Inefficient splicing events are not random since 32% are conserved from mice to humans. Two types of affected introns have been defined: nuclear-detained introns (DI), awaiting splicing completion or degradation, and retained introns (RI) subjected to <u>n</u>onsense <u>m</u>ediated <u>d</u>ecay (NMD). Whilst intron features influence retention events, the molecular mechanisms involved remain unclear (Boutz et al., 2015).

Following splicing, the EJC is deposited 24 nucleotides (nts) upstream of exon-exon junctions (Hauer et al., 2016; Le Hir et al., 2000) as a core of four proteins: the RNA helicase eIF4A3 which anchors the complex on RNA, the heterodimer Rbm8A-Magoh which locks eIF4A3 in its RNA-bound conformation, and Casc3 which binds RNA and stabilizes the complex (Andersen et al., 2006). The EJC also associates dynamically with peripheral proteins mediating its functions in splicing regulation, mRNA translation and stability (Boehm and Gehring, 2016).

As RNAPII transcribes the poly(A) site (pA site), the <u>c</u>leavage and <u>p</u>oly<u>a</u>denylation <u>c</u>omplex (CPAC) assembles on the pre-RNA to mature its 3’ end. CPA is important for transcription termination, mRNA export, stability and translation (Eckmann et al., 2011; Fong et al., 2015), and is in part regulated through modulation of cleavage site usage (also termed <u>a</u>lternative <u>p</u>oly<u>a</u>denylation, APA). APA impacts 70% of human mRNAs (Derti et al., 2012) and regulates cellular proliferation, tumorigenicity, and synaptic plasticity (Tian and Manley, 2017). Various human APA *trans*-regulators have been reported, some directly involved in CPA (Cpsf5, Cpsf6, Cstf2) (Martin et al., 2012; Masamha et al., 2014; Yao et al., 2012; Zhu et al., 2018)1, others linked to splicing (hnRNPC, Tardbp, U2af2, U1 snRNP) (Gruber et al., 2016; Kaida et al., 2010; Millevoi et al., 2006; Rot et al., 2017), and Thoc5, which plays a role in mRNA export (Katahira et al., 2013).

A major pathway for nuclear mRNA export uses the TREX complex (Strässer et al., 2002) which contains a core THO sub-complex, the RNA helicase Ddx39b, and RNA export adaptors and co-adaptors that Ddx39b loads onto mRNAs (Heath et al., 2016). A major adaptor is Alyref (Stutz et al., 2000), though some shuttling SR proteins can also work as adaptors (Huang and Steitz, 2003). Several co-adaptors have been characterized: Chtop, Thoc5, Cpsf6 and Rbm15 (Chang et al., 2013; Katahira et al., 2009; Ruepp et al., 2009; Zolotukhin et al., 2009). These proteins together recruit the export receptor Nxf1 and stimulate its RNA binding activity, which promotes mRNA export (Chang et al., 2013; Hautbergue et al., 2008; Viphakone et al., 2015; 2012). The topology of TREX components on RNAs and the molecular mechanisms involved in their deposition are still unclear. It has been suggested that the EJC may serve as a binding platform for RNA export factors (Le Hir et al., 2001; Singh et al., 2012). Yet, these interactions were not seen in other studies which instead revealed a key role for the <u>c</u>ap <u>b</u>inding <u>c</u>omplex (CBC, containing Ncbp1 and Ncbp2) in recruiting TREX, consistent with the idea that mRNPs may be exported 5’ end first (Cheng et al., 2006; Chi et al., 2013). Thus, the relative contribution of the EJC and CBC to TREX deposition on the RNA *in vivo* remains unresolved.

Here, we address some of these outstanding questions by performing <u>i</u>ndividual nucleotide resolution UV-<u>c</u>ross<u>l</u>inking and <u>i</u>mmuno<u>p</u>recipitation (iCLIP) (Broughton and Pasquinelli, 2013; König et al., 2011; Lee and Ule, 2018) on the mRNA export factors Alyref, Chtop, and Nxf1. Our *in vivo* results suggest co-transcriptional recruitment of these proteins all along the RNA during splicing but before CPA, which grants them additional roles in gene expression. Alyref can bind inefficiently spliced introns and regulates their splicing, and Chtop binds last exons and participates in APA regulation. We establish the EJC’s involvement in recruiting the major adaptor Alyref to spliced and single exon mRNAs but also to poorly spliced introns *in vivo*. Our data reconcile earlier disparate results by showing that the CBC acts as a transient landing pad for Alyref which is then transferred during co-transcriptional splicing to sites adjacent to the EJC throughout the RNA.

## RESULTS

### mRNA export factors bind protein coding transcripts with specific deposition patterns

To gain more insight into the population of RNAs bound by mRNA export factors and their distribution along RNAs *in vivo*, we used iCLIP. We generated stable cell lines expressing near endogenous levels of FLAG-tagged Alyref, Chtop, and Nxf1 which recapitulated known interactions (Figure S1 and (Chang et al., 2013; Fanis et al., 2012)), allowing us to use the same anti-FLAG antibody and identical conditions to permit direct comparisons between iCLIP data sets. For all three proteins, most binding events occurred within long, spliced RNAs, with approximately equal enrichment for classes such as protein-coding, lincRNA and pseudogenes. In contrast we observed much lower enrichment on short non-coding RNAs, such as snRNAs and ribosomal RNA (Figures 1A, 1B). Overall, 91%, 90% and 84% of the expressed protein-coding transcriptome had at least one clip tag for Alyref, Chtop, and Nxf1 respectively and 82% had clip tags for all three proteins. The fact that we didn’t observe any clear RNA-binding motif for any of those proteins, even within specific genic regions such as <u>u</u>n<u>t</u>ranslated <u>r</u>egions (UTRs), is consistent with the broad binding potential observed, and may be related to the use of unstructured stretches of arginines to bind RNAs by all three proteins (Chang et al., 2013; Hautbergue et al., 2008).

Among the lncRNAs bound by these RNA export factors was XIST (Figure 1C), which is consistent with XIST’s interactome (Chu et al., 2015). As most lncRNAs are chromatin-enriched and not exported (Schlackow et al., 2016; Werner and Ruthenburg, 2015), we identified nuclear and cytoplasmic lncRNAs for HEK293T cells using compartment-specific RNA-seq data (Sultan et al., 2014) and examined Alyref, Chtop, and Nxf1 binding to these RNA populations. Whilst Alyref and Chtop were enriched at similar levels regardless of the lncRNAs location, Nxf1 enrichment was reduced on nuclear lncRNAs compared to cytoplasmic ones (Figure 1D). This may contribute to the nuclear retention of lncRNAs, as previously suggested for XIST (Cohen and Panning, 2007).

**Figure 1.**
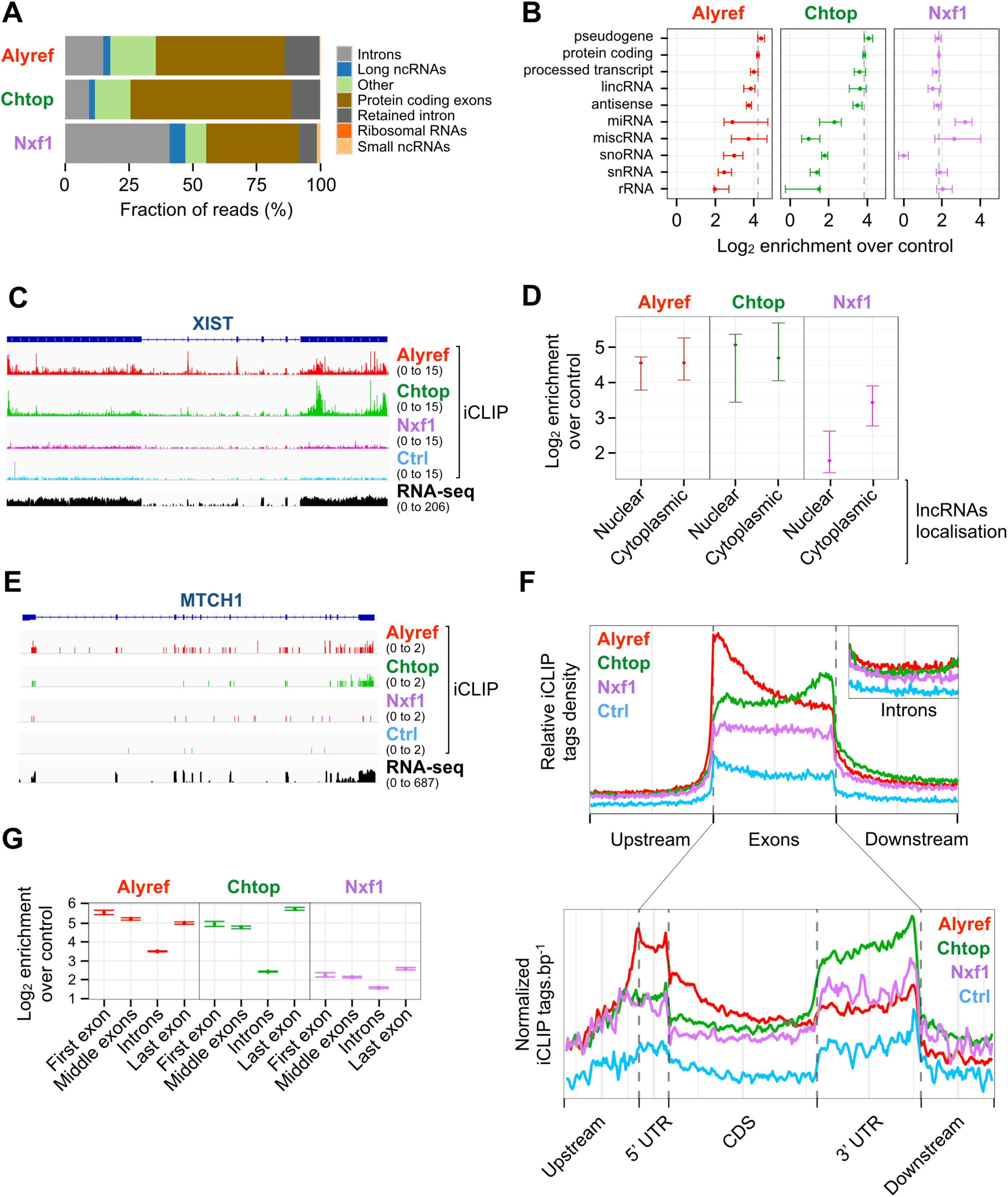
mRNA export factors bind protein coding transcripts with specific deposition patterns. **A.** Mapping context of binding clusters found in at least two replicates but not in control (Ctrl) iCLIP. The “Other” category contains miscRNA, NMD substrates and processed transcripts. **B.** Binding enrichment of RNA export factors over Ctrl iCLIP for the indicated gene biotypes. **C.** RNA export factor binding profiles over XIST lncRNA. **D.** Binding enrichments for nuclear and cytoplasmic lncRNAs. **E.** Example of an mRNA bound by RNA export factors. **F.** Deposition patterns of RNA export factors over the indicated genic regions. Data in the lower panel are normalised to RNA expression levels. **G.** Binding enrichment over Ctrl iCLIP for first/middle/last exons and introns. All binding enrichments are expressed as log_2_ crosslinks normalised to control. Error bars = 95% bootstrap confidence interval.

We found that the three proteins bifigurend along the whole transcript, but each with a specific profile. Alyref’s distribution displayed a 5’ bias due to a strong binding to first exons, consistent with its known connection with the CBC (Cheng et al., 2006), but interestingly it also exhibited binding events within the body of the RNA (Figures 1E, 1F, 1G). Chtop was also found on internal sites but its binding unexpectedly displayed a 3’ bias (Figures 1E, 1F) which corresponded to a strong enrichment on 3’ UTRs and last exons (Figures 1F, 1G). The RNA export receptor Nxf1 displayed a uniform distribution pattern across exons but was noticeably enriched on last exons (Figures 1F, 1G), a feature conserved in mouse (Müller-McNicoll et al., 2016). These specific profiles were absent on introns (Figure 1F). Of note, Alyref and Nxf1 iCLIP distribution profiles were almost identical to those of their respective yeast orthologs Yra1p and Mex67p obtained by PAR-CLIP (Baejen et al., 2014).

Together, these results showed the range of RNAs bound by these TREX components and revealed some intriguing deposition patterns.

### RNA export factors are loaded co-transcriptionally on spliced transcripts prior to 3’-end processing

The distributions of binding events detected within the body of RNAs prompted us to study the relationship between RNA processing status and recruitment of export factors. To this end, we computed a splicing index (SI, ratio of reads crossing internal exon-exon junctions to those mapping to exon-intron junctions) and a processing index (PI) (as described in (Baejen et al., 2014)). We found that these three TREX components had an SI >1 and a PI >2, indicating that they are mostly recruited to spliced but uncleaved RNAs *in vivo* (Figure 2A, 2B). Furthermore, while Alyref, Chtop, and Nxf1 are known nuclear proteins, we found by cellular fractionation that all three export factors are enriched in the chromatin fraction together with the TREX subunit Ddx39 (Figure 2C).

**Figure 2.**
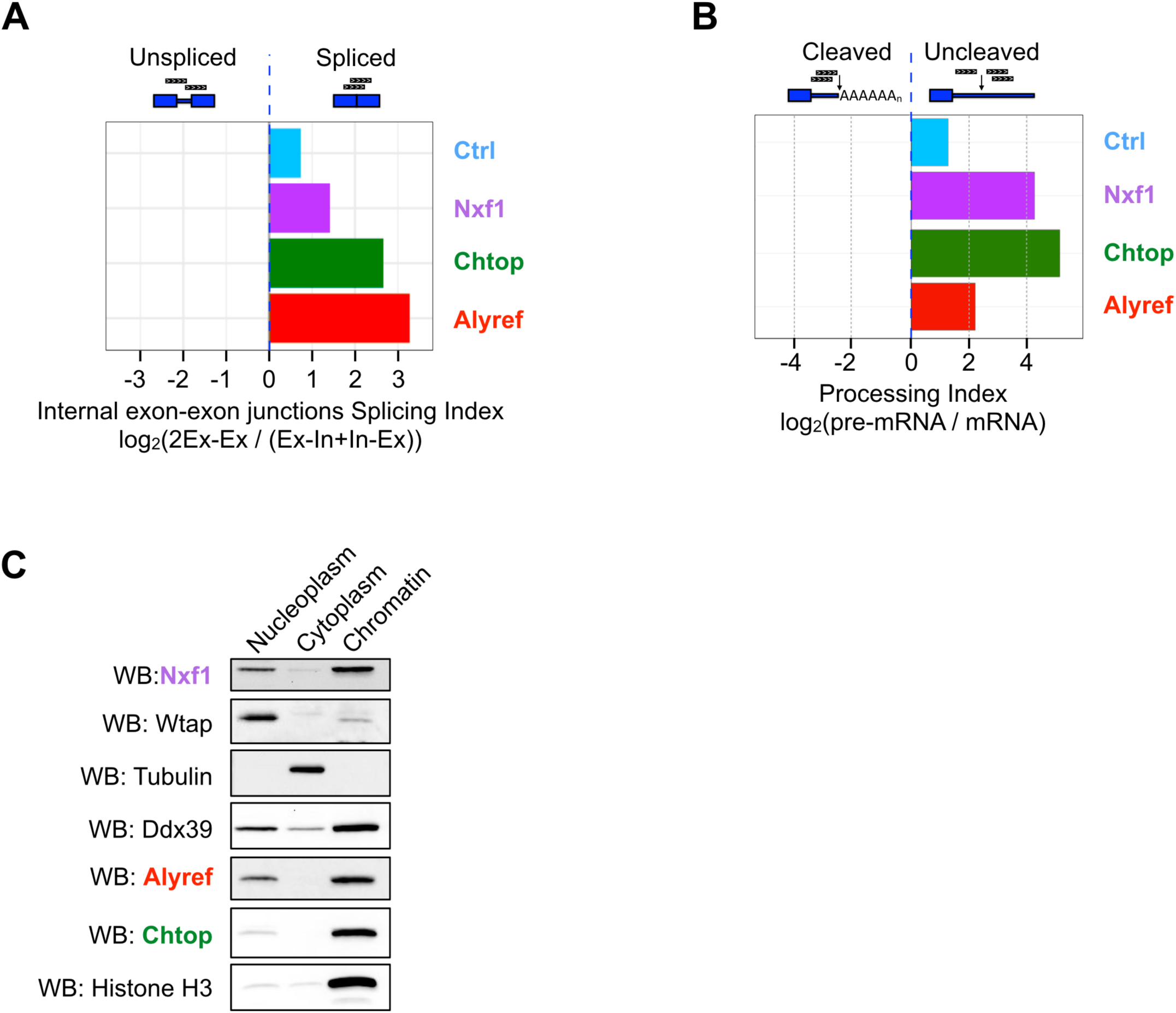
RNA export factors are loaded co-transcriptionally on spliced transcripts prior to 3’-end processing. **A.** Splicing indices are calculated as log_2_ ratio of spliced to unspliced reads at exon-exon junctions and indicate a binding preference for spliced versus unspliced RNAs for the designated factors. **B.** Processing indices are calculated as log_2_ ratio of iCLIP read depths for processed and unprocessed RNAs. **C.** Western blot (WB) analysis of 293T cells subcellular fractions using the indicated antibodies.

Since splicing is mostly co-transcriptional in human cells (Tilgner et al., 2012), together these results suggest that RNA export factors are recruited during transcription to the body of the RNA by the splicing process but before completion of 3’ end processing.

### Alyref binding to poorly spliced introns regulates their splicing *in vivo*

Consistent with co-transcriptional recruitment, we also detected some binding of RNA export factors to introns (Figure 1A). A known example of this is Nxf1 binding to an RNA secondary structure contained within the intron 10 of its own pre-mRNA (Li et al., 2006), which our data recapitulated (Figure S2). To look more closely at intronic binding events, we defined three classes of introns in HEK293T cells (see methods and (Boutz et al., 2015)): retained introns (RI), detained introns (DI), and finally introns not falling into these first two groups. Whilst the well-known splicing factor Ptbp1 was mainly enriched on regular introns, export factors were especially enriched on inefficiently spliced introns such as DI and RI, with the adaptor Alyref displaying the strongest enrichments (Figure 3A). Some examples of Alyref intronic binding are displayed in Figure 3B.

**Figure 3.**
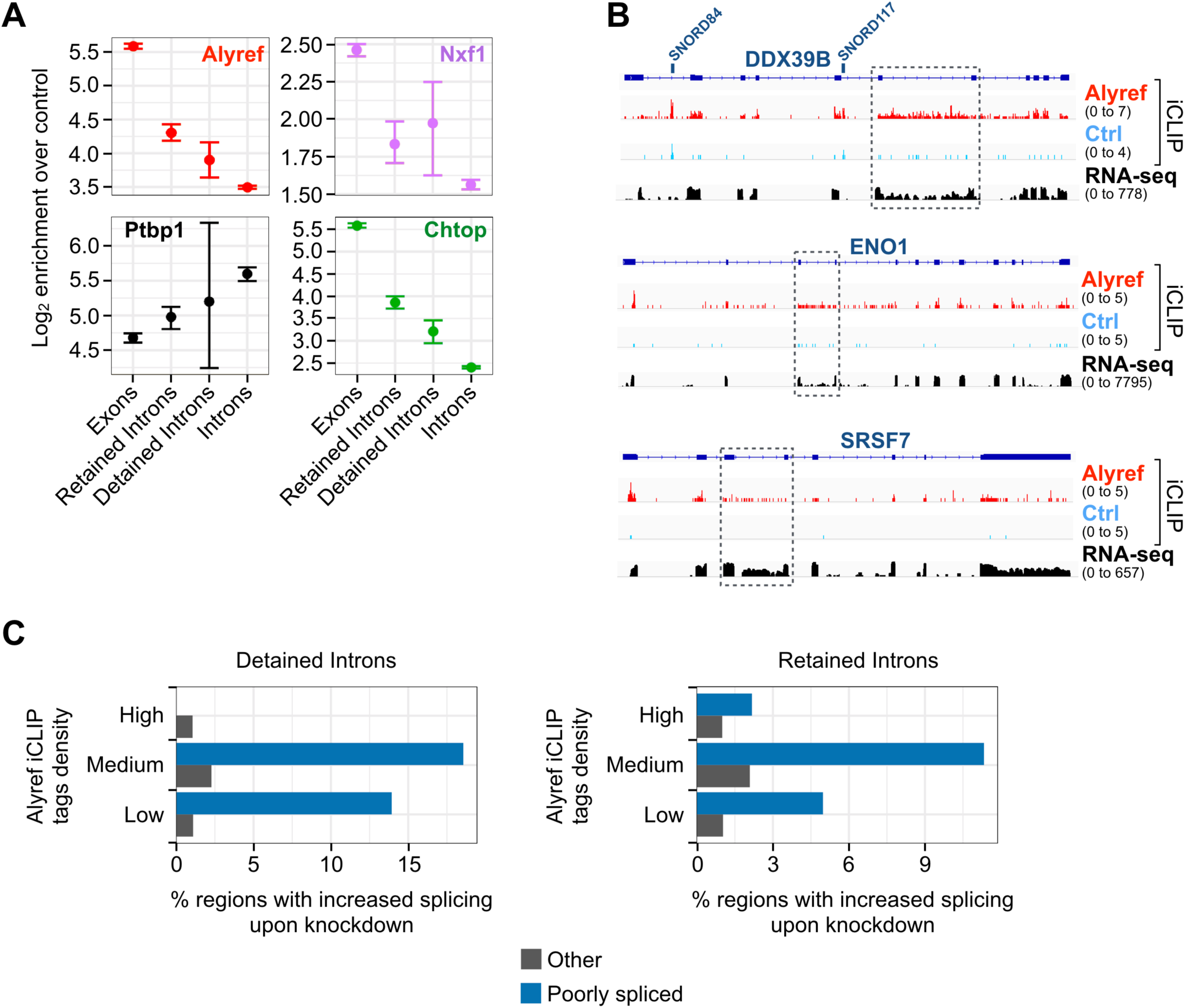
Alyref binding to poorly spliced introns regulates their splicing *in vivo*. **A.** Binding enrichment of RNA export factors and splicing factor Ptbp1 for the indicated RNA regions, calculated and displayed as in Figure 1. **B.** Examples of transcripts showing Alyref CLIP tags within introns that are normally inefficiently spliced (dashed boxes). **C.** Fraction of retained/detained introns and other transcript regions called as differentially included by DEXSeq (FDR 10%, Fold Change > 1.5) broken down by Alyref binding status.

To determine whether the binding of Alyref to these poorly spliced introns was functionally relevant, we investigated the effect of Alyref knockdown on RI and DI splicing using nuclear mRNA-seq data from HEK293T cells (Stubbs and Conrad, 2015). By using DEXseq to compare changes in RNA-seq signal over each intron with changes in other areas of their host transcript, we found that up to 18% of DI and 12% of RI bound by Alyref were more efficiently spliced upon Alyref depletion compared to < 2% of other transcripts regions (Figures 3C). These results suggest that mRNA export factors can also bind poorly spliced introns, and influence splicing outcomes.

### Chtop participates in APA regulation *in vivo*

The co-transcriptional recruitment of RNA export factors suggested by our data was also particularly interesting for Chtop for several reasons. Firstly, its strong processing index (Figure 2B, PI > 4) clearly suggested loading before 3’ end cleavage. Secondly, when analysed in more detail, Chtop binding events on 3’ UTRs were not affected by gene length and occurred within 1.5 kb of the 3’ end of the gene (Figure 4A), which matches the average size of 3’ UTRs in humans. By analyzing Chtop binding as a function of exon length, we found it preferentially bound long exons, unlike Alyref (Figure 4B). This probably explains the binding of Chtop to 3’ UTRs and last exons, which are on average the longest exons in transcripts. This trend persisted even when 3’ UTRs were excluded from the analysis (Figure S3A). Thirdly, the 3’ bias observed in Chtop binding profile was absent on replication-dependent histone mRNAs (Figure 4C), which are not usually polyadenylated in metazoans (Romeo and Schümperli, 2016). Finally, Chtop binds to the NTF2-like domain of Nxf1 (Chang et al., 2013) and interestingly, all the other confirmed or putative co-adaptors known to bind to this domain have been implicated in APA regulation (Katahira et al., 2013; Ruepp et al., 2009) (Figure 4D).

**Figure 4.**
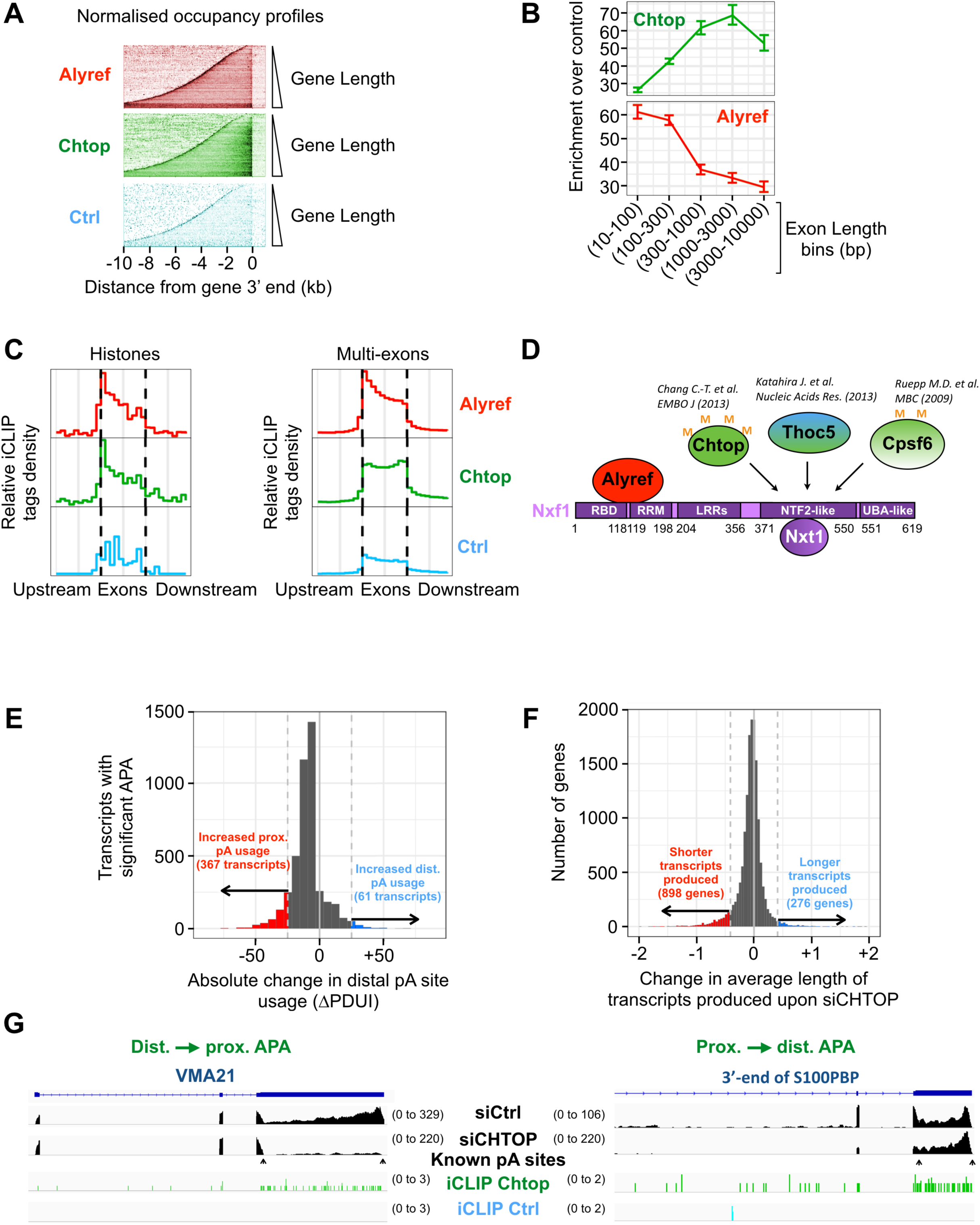
Chtop participates in poly(A) site choice *in vivo*. **A.** Occupancy profiles for the indicated factors and Ctrl iCLIP on transcripts produced from genes sorted by increasing length and aligned at their 3’ ends and normalised to nuclear RNA levels. **B.** Binding enrichment of Alyref and Chtop over Ctrl iCLIP as a function of increasing exon length. Error bars = 95% bootstrap confidence interval **C.** Deposition pattern of Alyref and Chtop over single exon histones and multi-exons RNAs using similar bins. **D.** Co-adaptors (in green-type colors) known to bind to Nxf1’s NTF2L domain. Thoc5 and Cpsf6 have been implicated in APA regulation in the indicated articles. **E.** Histogram showing the distribution of change in Percentage of Distal pA site Usage Index (ΔPDUI, (Xia et al., 2014)) for those transcripts found to have significant APA on Chtop knockdown (5% FDR) as detected by DaPars. Coloured regions show transcripts where |ΔPDUI| > 25%. **F.** Histogram of the log_2_ fold change in average effective gene length on Chtop knockdown, calculated by TxImport (Soneson et al., 2015) as an average of transcript lengths weighted by transcript usage. **G.** Examples of distal to proximal and proximal to distal APA changes observed upon Chtop depletion. RNA-seq tracks are shown in black. pA sites positions were from HEK293 A-seq (Martin et al., 2012). All replicates are shown in Figure S3C.

These results made us explore whether Chtop might regulate APA *in vivo*. To this end, we performed RNA-seq on poly(A)-selected total RNA from HEK293T cells depleted for Chtop. Of note, Chtop RNAi didn’t cause any change in the protein levels of the major APA regulator Cpsf6 or of Thoc5, also known to affect APA (Figure S3B). We observed 432 APA changes distributed as follows: a global reduction in distal pA site usage (Figure 4E, grey curve shifted to the left) with 367 distal to proximal and 61 proximal to distal significant changes. As an additional approach to analyze APA induced by Chtop depletion, we calculated transcript-specific expression levels and measured the per gene expression-weighted average transcript length. We observed that 892 genes produced shorter transcripts and 276 genes produced longer transcripts upon Chtop depletion (Figure 4F). Some examples of APA changes triggered by Chtop knockdown are presented in Figures 4G and S3C. As a reference, knockdown of Cpsf6 in HEK293T has been reported to lead to 775 significant APA changes (718 distal to proximal and 57 proximal to distal) ((Martin et al., 2012) data re-analyzed by (Zhu et al., 2018)). Correct pre-RNA 3’ end processing is intimately linked to transcription termination (Fong et al., 2015). Interestingly, we found that overexpression of Chtop leads to transcriptional read-through downstream of the CHTOP gene itself (Figure S3D).

Overall, these results suggest that the co-transcriptional recruitment of the RNA export factor Chtop to 3’ UTRs and last exons participates in defining the pattern of pA sites used *in vivo*.

### eIF4A3 is important for Alyref binding to RNA *in vivo* and mRNA export

The presence of RNA export factors on the body of RNAs prompted us to look at their distribution within those regions in more detail. To this end, we analysed their binding profiles in the vicinity of internal exon-exon junctions. As a point of reference, we re-analysed the EJC iCLIP data from (Hauer et al., 2016) and confirmed that this complex globally covers a region between −50 nts and −5 nts upstream of the exon-exon junction, peaking at −24 nts (Figure S4A). Remarkably, we observed an enrichment of Alyref in a region spanning over - 75 to −24 nts upstream of the exon-exon junction, with a peak at about −37 nts (Figure 5A). In contrast, this specific pattern was not observed for Chtop or Nxf1. Given the EJC’s known position (Figure S4A and Figure 5A brown arrowheads), this suggest that Alyref binds very close to the EJC within those internal regions of the RNA, illustrated with the schematic in Figure 5A.

**Figure 5.**
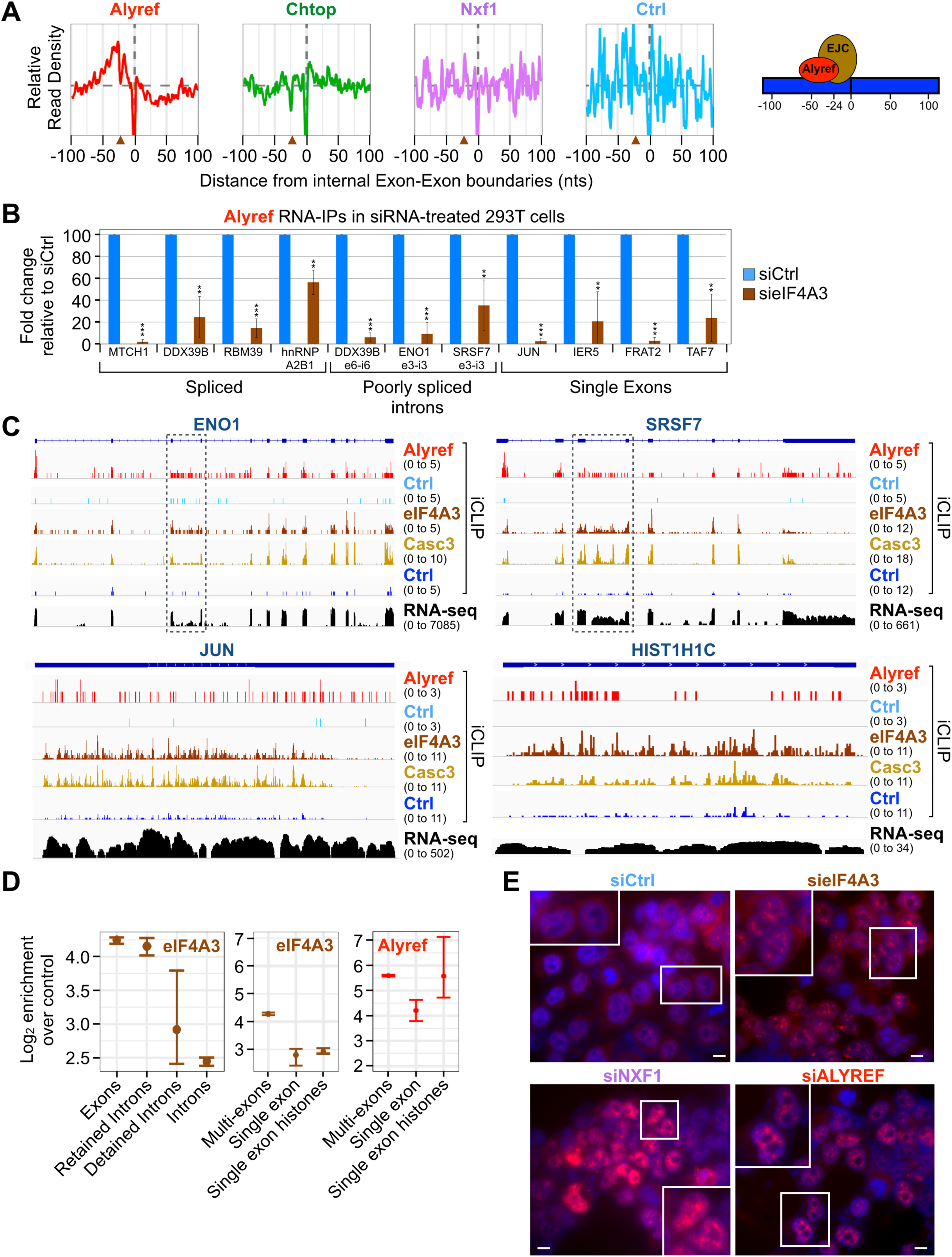
eIF4A3 is important for Alyref binding to a variety of RNAs and mRNA export. **A.** Average binding profile of RNA export factors at internal exon-exon junctions. The EJC’s position is inferred from Figure S4A and marked by brown arrowheads. **B.** Relative changes in Alyref RIP efficiency upon eIF4A3 RNAi in 293T cells. Means of three independent experiments +/- SD are shown. **P< 0.01, ***P < 0.001 (t-test). **C.** Examples of EJC bound to intronic (dashed boxes) or single exon RNAs. Also see Figure S4C. **D.** Binding enrichments of eIF4A3 and Alyref over Ctrl iCLIP on the indicated types of RNAs and RNA regions (as Figure 1). **E.** Oligo(dT) FISH on HeLa cells following RNAi for the indicated factors. Pictures were taken at the same exposure level. Some cells in a white box are shown at higher magnification. The scale bar corresponds to 10 µm.

The juxtaposition of Alyref and the EJC at exon-exon junctions and the earlier description of Alyref as a peripheral EJC component (Le Hir et al., 2000) prompted us to test the EJC’s importance for Alyref deposition at spliced junctions *in vivo*. Thus, we performed <u>R</u>NA-<u>IP</u>s (RIPs) on the endogenous Alyref protein using HEK293T cells depleted of eIF4A3, the EJC’s anchor. eIF4A3 RNAi efficiency was assessed by Western blot and RT-qPCR in the RIP samples (Figure S4B). Interestingly, eIF4A3 RNAi led not only to a clear reduction in the amount of spliced RNAs immunoprecipitated with Alyref, but also unspliced RNAs (Figure 5B), which Alyref can bind (Figure 3A). We initially included single exon transcripts as negative controls in these RIPs. To our surprise, eIF4A3 RNAi also severely reduced the amount of those RNAs immunoprecipitated with Alyref (Figure 5B). In agreement with these effects, we found the core EJC components eIF4A3 and Casc3 bound to these inefficiently spliced introns and to single exon transcripts (Figures 5C, S4C). Globally, these two core EJC subunits were particularly enriched on RI and DI over regular introns, albeit at a lower level than on exons, especially in the case of Casc3, and they were also enriched on histones and non-histones single exon RNAs, but less than on multi-exons (Figures 5D, S5A). In light of these results, we decided to test if eIF4A3 plays a role in mRNA export in human cells. eIF4A3 RNAi led to a moderate nuclear accumulation of poly(A)^+^-RNAs as assessed by oligo(dT) FISH, compared to the effects caused by depleting the export adaptor Alyref or the export receptor Nxf1 (Figures 5E, S5B). The strength of this phenotype was similar to that reported for Acinus, another EJC component (Chi et al., 2014). Additionally, eIF4A3 knockdown also led to an increase in the nucleo-cytoplasmic ratio of both single exon and spliced mRNAs (Figure S5C), consistent with a defect in nuclear export.

Together, these results show that the EJC is important for Alyref deposition onto an unexpected variety of transcripts and for the export of single-exon and spliced RNAs.

### CBC acts as a transient landing pad for Alyref

As the majority of exon junctions lie within the body of mRNAs, the EJC is mostly present on ORFs or “middle exons” (Hauer et al., 2016), a result that we confirmed (Figure S5D). Therefore, the significant role played by the EJC in recruiting Alyref that we had uncovered was puzzling given the Alyref 5’ enrichment (Figures 1F, 1G). Indeed, since the 5’ cap of eukaryotic mRNAs is added soon after transcription initiation, Alyref 5’ bias and its known association with the CBC (Cheng et al., 2006) would suggest that Alyref could bind early to the 5’ end, possibly before the first intron of the transcript has been excised. However, we found by RIPs that Alyref preferentially bound to 5’ ends that had already undergone splicing (Figure 6A). To analyse this phenomenon more globally, we computed a splicing index restricted to the first exon-exon junctions. We found that Alyref preferentially binds to spliced RNAs, even at the 5’ end (Figure 6B). These results suggested that Alyref could be tethered at the 5’ end of RNAs by the CBC but that it still required splicing for binding, as previously proposed (Cheng et al., 2006).

**Figure 6.**
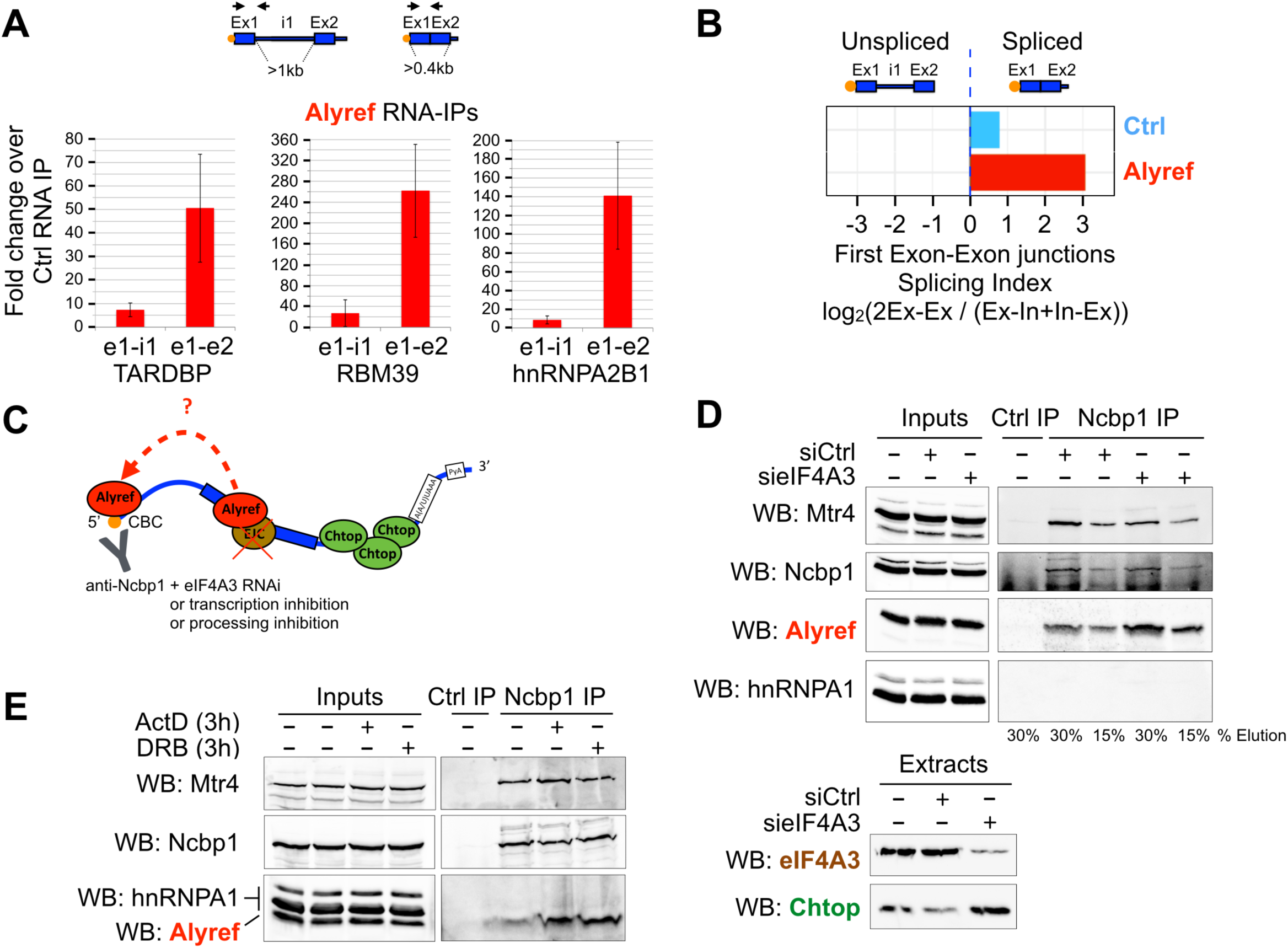
CBC acts as a transient landing pad for Alyref. **A.** Analysis of the influence of splicing on endogenous Alyref binding to the 5’ end of selected transcripts by RIP. Means of three independent experiments +/- SD are shown. **B.** Splicing indices at first exon-exon junctions computed for Alyref and iCLIP Ctrl as in Figure 2. **C.** Experimental strategy used in panels D and E. **D.** Influence of eIF4A3 RNAi on Alyref co-IP with the CBC subunit Ncbp1 analysed by western blot. hnRNPA1 is a negative control. Mtr4 is a known CBC interactor. Efficient depletion of eIF4A3 is shown in the lower panel. **E.** Influence of transcription inhibition using ActD or DRB on Alyref co-IP with Ncbp1 analysed by western blot.

In an attempt to explain these discrepancies, we hypothesized that after a pre-requisite initial recruitment of Alyref by the EJC, the potential spatial proximity between EJCs and the CBC induced by EJC-mediated compaction of the mRNP (Singh et al., 2012), might allow a transfer of Alyref to the CBC. To test this, we immunoprecipitated CBC and looked at the levels of Alyref co-purifying in an eIF4A3 RNAi context (Figure 6C). If this hypothesis was true, then eIF4A3 RNAi should result in a reduced amount of Alyref bound to CBC. Surprisingly, we observed the opposite: Alyref specifically accumulated on the CBC upon eIF4A3 RNAi (Figure 6D).

We then performed CBC IPs after transcription inhibition by actinomycin D or DRB which also led to accumulation of Alyref on CBC (Figures 6C, 6E). Interestingly, the same phenomenon was observed upon splicing inhibition with pladienolide B (Kotake et al., 2007), whereas treatment with cordycepin, an inhibitor of polyadenylation (Kondrashov et al., 2012), had no effect (Figures 6C, S6A). To confirm that the two drugs were active in this experiment, we performed RT-qPCR on total RNA extracted from a fraction of the IP inputs. Splicing was inhibited, as shown by an increase in some unspliced RNAs, and cordycepin treatment led to an expected increase of the PAXT complex RNA target SNHG19, as observed previously (Meola et al., 2016) (Figure S6B).

Overall, these *in vivo* results show that interfering with co-transcriptional recruitment of Alyref can lead to its accumulation on the CBC. In combination with our iCLIP data, this suggest that, *in vivo*, Alyref binds to the CBC initially but in a transient manner, before being subsequently deposited near the EJC on the RNA.

## DISCUSSION

Based on the data presented here, we propose a model for the co-transcriptional recruitment of RNA export factors and their influence on gene expression *in vivo* (Figure 7), which unifies two long-standing disparate views: a CBC-only model or an EJC-centric model. As the RNA exits RNAPII, the CBC elicits early recruitment of mRNA export factors at the 5’ end. As components of the spliceosome rapidly scan the RNA, this may suffice to deposit the EJC and mRNA export factors on single exon transcripts, but at a lower efficiency than on intron-containing RNAs (Figures 5D, S5A). On multi-exon transcripts, splicing deposits the EJC. Alyref then transfers from the CBC to a site upstream of the EJC at all exon-exon junctions along the RNA. Intron retention can also trigger atypical binding of the EJC and Alyref, which in turn regulates this phenomenon. As the last exon is produced by RNAPII, Chtop presence increases and it participates in APA regulation. Ultimately, Alyref and Chtop allow Nxf1 to join the RNP and stimulate nuclear export.

**Figure 7.**
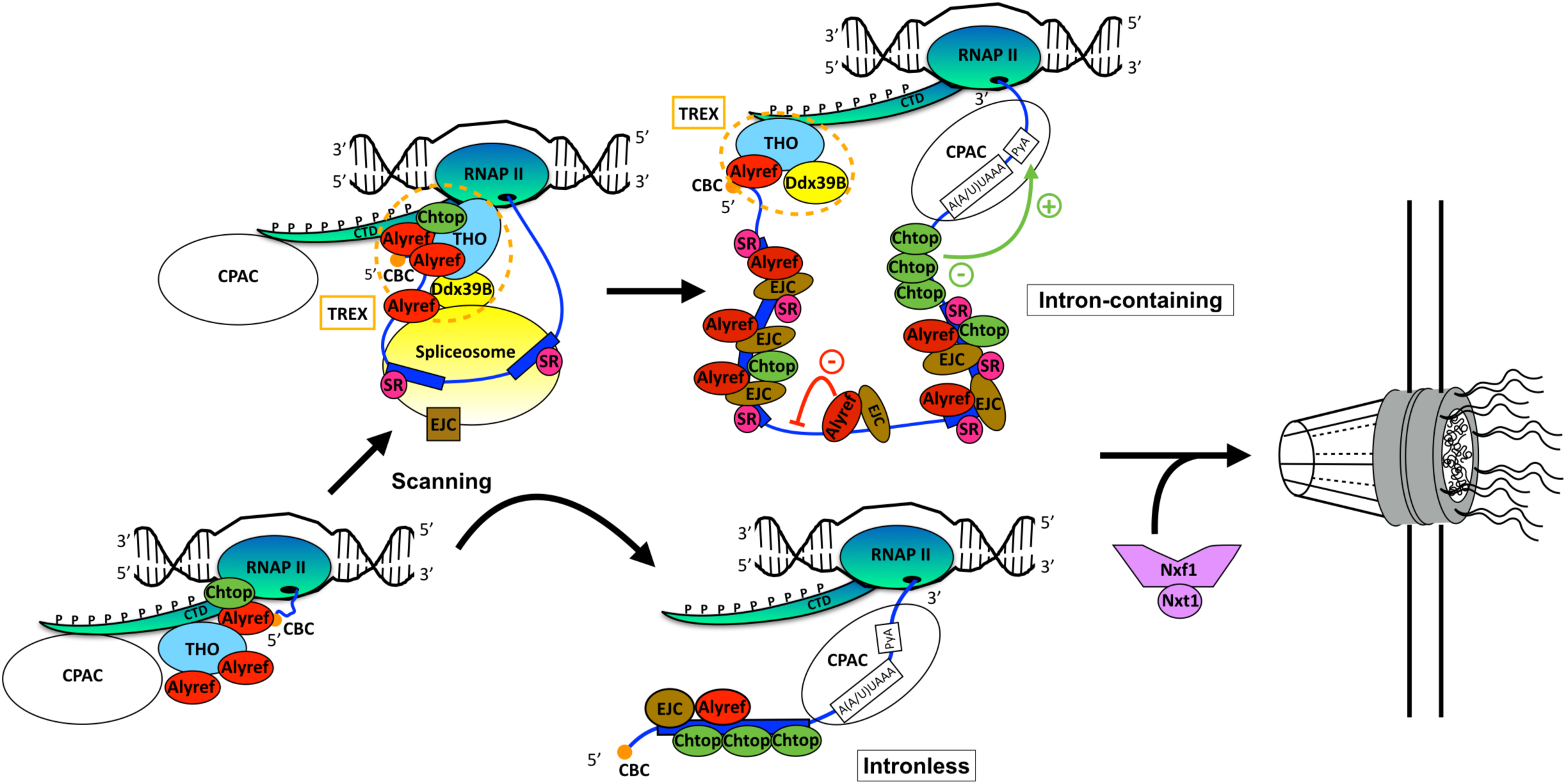
A model for the co-transcriptional recruitment of mRNA export factors and its effect on gene expression. Proposed model for the co-transcriptional recruitment of RNA export factors and their influence on gene expression *in vivo*.

Since TREX and the EJC can be loaded on RNAs via splicing but independently of transcription *in vitro* (Cheng et al., 2006), we cannot rule out that some Alyref may be loaded post-transcriptionally. However, this probably accounts for a minority of events, as splicing is mostly co-transcriptional in human cells. Although our data could also reflect an influence of the CBC in the splicing of the first intron (Ohno et al., 1987), our model is supported by the following recent lines of evidence. Alyref associates with spliced RNAs in a cap- and EJC-dependent manner *in vitro*, but a simultaneous binding of Alyref to both CBC and EJC seems unlikely *in vivo* ((Gromadzka et al., 2016) and see below). Moreover, the CBC isn’t present in the EJC’s interactome (Singh et al., 2012). Consistent with Alyref being only transiently associated with CBC *in vivo*, Alyref isn’t strongly enriched in CBC interactome studies at steady state, unlike Mtr4/Skivl2, or THO components (Andersen et al., 2013; Gebhardt et al., 2015; Pabis et al., 2013). Furthermore, we found Alyref bound all along the RNA (Figures 1F, 1G) and depletion of the CBC subunit Ncbp1 decreases Alyref binding not only at the 5’ end but also throughout the body of the RNA (Shi et al., 2017). Moreover, the CBC now appears to be critical for poly(A)^+^ RNA export in human cells contrary to previous studies (Gebhardt et al., 2015). The repeated loading of Alyref from CBC onto internal sites juxtaposed with EJCs would ensure that the CBC remains loaded with Alyref throughout mRNP maturation. This is likely to be important in mRNP quality control as Alyref competes with Mtr4 for CBC interaction and this competition determines whether an mRNA is exported or, if defective, degraded by the exosome (Fan et al., 2017).

The co-transcriptional nature of the recruitment of RNA export factors has several implications. Firstly, it shows that Alyref binding to poorly spliced introns participates in regulating their excision transcriptome-wide (Figure 3). Although initially considered as a biological defect, intron retention is now regarded as part of crucial regulatory programs activated during physiological or pathological gene expression, such as cellular differentiation (Llorian et al., 2016), response to neuronal activity (Mauger et al., 2016), and tumor suppressor inactivation in cancer (Jung et al., 2015). Our findings suggest that canonical mRNA export factors have the potential to play a role in these processes.

Secondly, co-transcriptional recruitment raises questions regarding the various signals required to release the RNP from the chromatin for nuclear export. Nxf1’s presence is likely to be one of those signals, given its reduced levels on nuclear-retained lncRNAs (Figure 1D) and the export competence that it grants even to unspliced transcripts (Braun et al., 2001; Li et al., 2006). Yet, the binding of RNA export factors to poorly spliced introns (Figure 3B) is overall clearly insufficient since only 10% of all RI-containing transcripts artificially induced by splicing inhibition reach the cytoplasm (Yoshimoto et al., 2016). Another important signal is probably the RNA’s maturation state since inefficient processing impairs release from the site of transcription and leads to degradation by the nuclear exosome (Custodio et al., 1999; Schlackow et al., 2016). Interestingly, Alyref loading at the 5’ end seems to help RNAs evade nuclear degradation by the exosome (Fan et al., 2017). In mammals, there is growing evidence that splicing more than 3’ end formation is responsible for the nuclear retention of transcripts and their release for export. Indeed, poly(A)^+^ RNAs can be found at transcription sites after transcription termination, and splicing completion can delay the release of fully 3’ end-formed RNAs from the chromatin (Brody et al., 2011; Mauger et al., 2016; Pandya-Jones et al., 2013). Moreover, lncRNAs are mainly chromatin-associated (Werner and Ruthenburg, 2015) and splicing more than 3’ end formation seems to define their poor processing state leading to their degradation by the nuclear exosome (Schlackow et al., 2016). In line with this, we show that the splicing process is key to allow co-transcriptional recruitment of export factors (Figure 2) and that unspliced species are indeed recruiting export factors less efficiently (Figure 3A, 5D). They are also consistent with the roles played by splicing and TREX components in releasing spliced RNAs from nuclear speckles and stimulating mRNA export (Dias et al., 2010).

Thirdly, our data on Chtop participating in APA regulation (Figure 4) support the view that the co-transcriptional loading of RNA export factors can influence 3’ end processing in human cells. Since Cpsf6 is a master regulator of APA in the cell, we looked at the set of transcripts displaying APA changes in both siCPSF6 (Martin et al., 2012) and siCHTOP conditions. It was restricted to a few transcripts (21), revealing the complex interplay taking place on last exons to establish the pattern of pA sites used in the cell. Interestingly, Nxf1 was up-regulated in response to Chtop depletion (Figure S3B) and Nxf1 knockdown triggers hyperadenylation of reporter transcripts (Qu et al., 2009). These results reinforce the functional connection that we have previously uncovered between Nxf1 and Chtop (Chang et al., 2013). Consistent with an involvement of Chtop in 3’ end processing regulation, its overexpression specifically led to transcriptional read-through downstream of its own gene (Figure S3D). Perhaps this constitutes another mechanism used by Chtop to auto-regulate its expression (Figures S1, S3D and (Izumikawa et al., 2016)).

Although the EJC wasn’t originally implicated in RNA export, those data were either obtained *in vitro* (Cheng et al., 2006), or before the discovery of the EJC’s anchor eIF4A3. Interestingly enough, only the combined depletion of all EJC subunits known at the time triggered a partial block of export, and the co-depletion of Alyref and these EJC proteins is lethal in Drosophila cells (Gatfield and Izaurralde, 2002). Our study confirms a key functional link between the EJC and Alyref. In agreement, Alyref binding and enrichment patterns mirrored those of the EJC, whereas Chtop and Nxf1 were more alike (Figures 5A and S4A, and Figures 4B, S3A, S5E). Also, depletion of either Alyref or eIF4A3 increases the levels of Chtop. Both cases could be a cellular response to the loss of Alyref on the RNA, since Alyref works with Chtop to mediate TREX function (Figure 6D and (Chang et al., 2013)). Since endogenous Alyref interacts poorly with EJC components (Cheng et al., 2006; Chi et al., 2013) and isn’t strongly enriched in the EJC interactome (Singh et al., 2012), its proximity to the EJC is perhaps achieved through RNA-enhanced protein-protein interactions. Interestingly, a recent study found that mutating a conserved motif in Alyref abrogates the interaction of FLAG-tagged Alyref with both the CBC and the EJC, suggestive of a mutually exclusive site (Gromadzka et al., 2016). We speculate that this motif might allow Alyref to transfer from the CBC next to the EJC.

Surprisingly, our work has revealed the presence of the EJC on intronless RNAs. Our results are consistent with the ability of EJC subunits to bind nascent transcripts independently of pre-mRNA splicing in Drosophila (Choudhury et al., 2016). Moreover, eIF4A3 enhances translation of intronless mRNAs by binding to their 5’ end ((Choe et al., 2014), Figure S5F). The recruitment of splicing factors to intronless mRNAs is not unprecedented and could result from a “scanning” mechanism of the pre-mRNA by spliceosome components (Änkö et al., 2012; Brody et al., 2011). The whole EJC is associated with the 5’ exon prior to exon ligation and eIF4A3 is the only EJC core protein to directly interact with spliceosome subunits (Zhang et al., 2017). Also, eIF4A3 doesn’t seem to recognize exons as well as Casc3 (Figures 5D, S5A), nor their positions within the transcript (Figure S5D). Therefore, it could be involved in such a scanning process. Intronless RNAs are notoriously poorly exported and are also subject to greater exosome-mediated degradation than regular mRNAs (Dias et al., 2010; Fan et al., 2017). Interestingly, Alyref and the EJC were both more enriched on multi-exons than single-exon RNAs (Figures 5D, S5A), whereas Chtop and Nxf1 were not discriminative (Figure S5A). Given that the major mRNA export adaptor Alyref seems to help RNAs evade nuclear degradation by the exosome (Fan et al., 2017), it is plausible that Alyref’s loading might be a rate-limiting step for the efficient export of intronless RNAs. In contrast, the single exon histone mRNAs, which bind TREX subunits (but not the EJC) just as strongly as multi-exon RNAs, may have evolved more efficient ways to recruit RNA export factors (Figures 5D, S5A).

Overall, our study highlights an important role for both the CBC and the EJC in the nuclear export of RNAs in human cells and shows that, upstream of their role in stimulating export, the binding of RNA export factors to their RNA targets plays an important a role in shaping the transcriptome by influencing splicing decisions and 3’ end processing.

## MATERIAL & METHODS

### CONTACT FOR REAGENTS

Stuart A. Wilson

### EXPERIMENTAL METHODS

Cell lines and tissue culture conditions

HEK293T, HeLa, cell lines were maintained in Dulbecco’s modified Eagle medium (DMEM) with 10% foetal bovine serum (FBS). FlpIn-293 cells expressing the FLAG-tagged proteins were generated as described previously (Hautbergue et al., 2009) and maintained in DMEM with 10% FBS, 15 µg/mL Blasticidin, and 0.1 mg/mL Hygromycin.

#### siRNA transfections

Cells were transfected on the day of seeding with 7.5 nM of siRNAs (30 nM for HeLa cells) using RNAiMAX (Life technologies) according to the manufacturer’s instructions. The transfection was repeated at 48 hours and the cells were harvested between 68 and 72 hours after the first transfection. The siRNAs used were: siCtrl 5’-CACCGUGAAGCUGAAGGUG-3’ (Viphakone et al, 2015), sieIF4A3 5’-AGCCACCUUCAGUAUCUCA-3’, siChtop 5’-GACAACCAAUUGGAUGCAUAU-3’, siNXF1 5’-UGAGCAUGAUUCAGAGCAA-3’.

#### Fluorescent In Situ Hybridisation (FISH)

Performed as described in (Viphakone et al., 2012).

<u>i</u>ndividual nucleotide resolution <u>C</u>ross<u>L</u>inking and <u>I</u>mmuno<u>P</u>recipitation (iCLIP)

Near endogenous expression of the FLAG-tagged proteins was induced for 48h in the stable cell lines by addition of tetracycline at 5 ng/mL for Nxf1, and 10 ng/mL for Chtop, and Alyref. iCLIP was then performed exactly as described previously (König et al., 2011) but using 10 µg of anti-FLAG antibodies (Sigma) and performing the high salt washes of the immunoprecipitation part of the protocol at room temperature (22°C) on a rotating wheel for 5 minutes per wash. Additionally, the following modifications from (Broughton et al., 2013) were used. All nucleic acid pellets were air dry for a maximum of 10 minutes to ease resuspension. Nucleic acids centrifugations after each overnight precipitation were done at 4°C, 16000 × g for 20 minutes and 5 minutes for the EtOH 80% washes. During the cDNA gel purification step, each gel piece was transferred to a 0.5 mL microcentrifuge tube pierced with 3-4 holes made with a 19-gauge needle. The gel-containing 0.5 mL tube was itself inserted inside a 1.5 mL microfuge tube and the assembly was centrifuged at 16000 × g for 2 minutes to efficiently shred the gel piece. After circularisation of the cDNAs, the Circligase was inactivated by heating the samples for 15 minutes at 80°C. To linearize the cDNAs, the standard iCLIP cut_oligo was replaced by a cut_oligo bearing a 3’ dideoxycytosine instead of four adenosines. Libraries for each protein of interest, as well as libraries from cells expressing only the FLAG tag were prepared in triplicate. Libraries were pooled, a spike-in of 10% PhiX DNA added and distributed across 4 lanes of Illumina HiSeq 2500 and 1 lane of Illumina MiSeq sequencing by the Centre for Genomic Research (Liverpool, UK).

#### Formaldehyde RNA Immuno-Precipitation (faRIP)

One 6-cm dish (or 2 × 6-cm dishes for siRNA treatments) was seeded per RIP condition with 300000 cells/dish. Protein-RNA complexes were crosslinked *in vivo* 48 hours later (or 68-72 hours later for siRNA treatments) by incubating the cells with 3 mL of PBS-Formaldehyde (0.1%). 100 µL of protein G-Dynabeads were prepared by initial washing with 3 × 1 mL RIP lysis buffer (50 mM HEPES-HCl pH 7.5, 150 mM NaCl, 10% glycerol 1% NP-40, 0.1% SDS, and 0.5% sodium deoxycholate) before being blocked and loaded with the relevant antibody (4 µg diluted in 0.3 mL of RIP lysis buffer + 1% BSA w/v final; control RIP antibody was anti-FLAG, Sigma) for 1 hour at room temperature. The beads were then washed with 3 × 1 mL RIP lysis buffer and left on ice until further use. Each cell pellet was lysed in 400 µL RIP lysis buffer supplemented with 1 mM DTT, protease inhibitors (SigmaFAST, Sigma), 2 µL of RNase inhibitors (Ribosafe, Bioline) and 2 µL/mL of Turbo DNase (Ambion). Samples were then sonicated using a Bioruptor (High, 8 × [30s-ON/30s-OFF]) to generate fragments of ~ 350 nts, and cleared by centrifugation (16100 × g, 10 minutes, 4°C). 300 µL of each sample were incubated with the prepared Dynabeads for 2 hours at 4°C and 30 µL of lysate were kept as an input (10%). Following incubation, the beads were washed with 2 × 1 mL RIP lysis buffer, 2 × 1 mL high salt RIP lysis buffer (adjusted to 500 mM NaCl, 5 minutes each on ice), and again 2 × 1 mL RIP lysis buffer. Crosslinks reversal and elutions were performed by adjusting inputs and washed beads to 100 µL with reverse-crosslinking buffer (final concentrations: PBS 1X, 2% N-lauroyl sarcosine, 10 mM EDTA, 5 mM DTT, 1.9 mg/mL proteinase K (Roche)) and shaking them at 1100 rpm for 1 hour at 42°C followed by 1 hour at 55°C. The RNA content of the resulting eluates and inputs were extracted using TRIzol (Life Technologies), following the manufacturer’s instructions. All RNA samples were then DNase-treated (Turbo DNase, Ambion), phenol/chloroform extracted, ethanol precipitated, and resuspended in RNases-free water (Sigma). The whole content of RNAs obtained from the immunoprecipitations and inputs were used for cDNA synthesis and qPCR analysis.

#### Co-Immunoprecipitations (co-IPs)

HEK293T cells were seeded at 60% confluency in 10-cm dishes. Where indicated, the cells were treated with either actinomycin D (5 µg/mL, Sigma), or 5,6-dichlorobenzimidazole 1-ß-D-ribofuranoside (DRB, 100 µM, Sigma), or pladienolide B (1 µM, Santa Cruz Biotechnology, sc-391691, Kotake et al., 2007), or cordycepin (25 µM, Kondrashov et al., 2012), or DMSO (as mock-treated negative control) for 3 hours prior to harvesting. 4 µg of the indicated antibodies (or anti-FLAG for control IPs) were incubated with 100 µL of protein G Dynabeads for 1 hour at room temperature in 300 µL of IP lysis buffer (50 mM HEPES-NaOH pH 7.5, 100 mM NaCl, 1 mM EDTA pH 8, 0.1% Triton X-100, 10% Glycerol) supplemented with 1% BSA. The cells were briefly washed in ice-cold PBS and lysed in ice-cold IP lysis buffer supplemented with 1 mM DTT, Turbo DNase (4 U/mL), RNase A (10 µg/mL) and protease inhibitors (SigmaFast). Equivalent amounts of cleared protein extracts were incubated with the prepared antibody-bound protein G Dynabeads for 2 hours, washed three times with 1 mL of IP lysis buffer containing RNase A (10 µg/mL) and eluted by an acid shock using 1 M Arginine-HCl (pH 3.5) and neutralisation to pH 7.5 using 1.5 M Tris-HCl pH 8.8. 0.1-0.2% of inputs and 16-30% of eluates were subsequently analysed by SDS-PAGE and Western blotting using the indicated antibodies.

#### RNA-seq on siCtrl- and siCHTOP-treated cells

600 000 cells were seeded in a 10-cm dish and subjected to siRNA-mediated knockdown (see above). Total RNA extraction was performed using TRIzol following the manufacturer’s instructions. Samples were resuspended in 50 µL of RNase-free water (Sigma), DNase-treated with 4 U of Turbo DNase (Ambion) for 1 hour at 37°C, phenol-extracted, ethanol-precipitated overnight at −20°C and resuspended in 40 µL of RNase-free water. mRNA enrichment, cDNA generation, strand-specific library preparation and sequencing were performed using standard Illumina protocols by Novogene (Beijing, China). At least 20 millions read pairs were generated.

#### Cellular fractionation and RNA/Protein extraction

Nuclear and cytoplasmic proteins and RNAs were extracted from cells as described previously (Stubbs et al., 2015) with the following two modifications: firstly Ribosafe (0.1 U/mL) was used in place of RNasin (0.04 U/mL), and secondly, after cell lysis, the nuclei were additionally washed twice with “Buffer I” (10 mM Tris pH 8.0, 0.32 M Sucrose, 3 mM CaCl_2_, 2 mM MgCl_2_, 0.1 mM EDTA, 1 mM DTT, 10% glycerol). To extract nucleoplasmic and chromatin-associated proteins, the purified nuclei were resuspended in 350 µl of NRB buffer (20 mM HEPES pH 7.5, 50% Glycerol, 75 mM NaCl, 1 mM DTT, protease inhibitors SigmaFast), and then an equal volume of NUN buffer (20 mM HEPES, 300 mM NaCl, 1 M Urea, 1% NP-40 Substitute, 10 mM MgCl_2_, 1 mM DTT) was added and incubated 5 minutes on ice, then centrifuged (1,200 × g, 5 minutes, 4°C). The supernatant was transferred to another tube and corresponded to the soluble nucleoplasmic extract. The crude chromatin pellet was resuspended in 1 ml of Buffer A (10 mM HEPES pH 7.5, 10 mM KCl, 10% glycerol, 4 mM MgCl_2_, 1 mM DTT, protease inhibitors SigmaFast) to wash, transferred to another 1.5 mL tube, and centrifuged (1,200 × g, 5 minutes, 4°C). The resulting purified chromatin pellets were resuspended in 100 µL of RIPA buffer (50 mM HEPES pH 7.5, 150 mM NaCl, 1 mM DTT, 10% glycerol, 1% NP-40, 0.1% SDS, 0.5% sodium deoxycholate + protease inhibitors SigmaFast) and treated with 500 U of Benzonase at room temperature for 45 minutes to release the chromatin-associated proteins. The digested chromatin was cleared by centrifugation (16100 × g, 10 minutes, 4°C) and the supernatant was analysed by SDS-PAGE and Western blot with the indicated antibodies.

### BIOINFORMATICS METHODS

#### Other datasets used

eIF4A3 and Casc3 iCLIP data were downloaded from ENA accession ERA551949 (Hauer et al., 2016). Total nuclear and cytoplasmic ribosome-depleted RNA-seq from HEK293 cells was downloaded from ENA accession PRJEB4197 (Sultan et al., 2014). Nuclear mRNA-seq from HEK293 was downloaded from GEO accession GSE111878 (manuscript submitted). HEK293 A-seq was downloaded from GEO accession GSM909242 (Martin et al., 2012). Nuclear and cytoplasmic mRNA-seq data from Control and Alyref knockdown HEK293 cells were the kind gift of Nicolas Conrad (Stubbs et al., 2015) and were provided as pre-mapped alignments.

#### Gene Sets

Throughout the study two gene sets were used. The ‘reference’ set was obtained from Ensembl 75. The ‘expressed’ set was derived using transcripts with mean TPM > 1 as measured by Salmon 0.8.2 (Patro et al., 2017) using total nuclear RNA-seq. We excluded genes on the mitochondrial chromosome. To annotate different parts of genes, we first divided genes from the reference set into the minimum number of non-overlapping chunks such that any isoform of the gene could be constructed from a combination of these chunks. This was performed using the ‘genes-to-unique-chunks’ mode of the ‘gtf2gtf’ tool from CGAT (Sims et al., 2014). For each chunk, we then counted the number of exons and introns it overlapped with from both the reference and expressed gene sets using ‘bedtools intersect’. We excluded any chunks that overlapped with more than one gene. We defined introns as chunks that overlapped with at least one intron from the expressed transcript set, but no exons from any isoform; and exons as chunks that overlapped with at least one exon from the expressed transcript set, but no introns from any isoform.

#### iCLIP data processing

iCLIP sequencing data was processed using pipeline_iCLIP distributed as part of iCLIPlib (http://www.github.com/sudlab/iCLIPlib). For data generated for this study, we removed spiked in phiX sequence, by mapping reads against the phiX genome using bowtie 1.1.2 (Langmead et al., 2009) with the settings ‘-v 2 --best --strata -a’. UMI sequences were extracted using UMI-Tools 0.5.3 (Smith et al., 2017) leaving the sample barcode on the read sequence. Reads were then simultaneously trimmed and demultiplex using Reaper from the Kraken tools (Davis et al., 2013). Only reads longer than 15 nts were retained. The remaining reads were then mapped against the hg19 genome sequence using STAR 2.4.2a (Dobin et al., 2013) and a junction database built from Ensembl 75. The options to STAR were ‘--outFilterMultimapNmax 1 --outFilterType BySJout --outFilterMismatchNoverLmax 0.2 --outFilterScoreMin 0.8 --alignSJDBoverhangMin 1’. Thus, multimapping reads, reads with greater than 20% mismatches or an alignment score less than 0.8 were discarded. Only splice junctions present in Ensembl 75 were allowed, but only an overhang of 1 nucleotide was required for splicing at an annotated junction. This prevents spuriously overhanging sequences from affecting calculations of splicing ratio (see below). Reads from the same sample sequenced on different lanes were then merged and PCR duplicates removed accounting for sequencing and PCR errors using UMI-Tools ‘dedup’. For each read the cross-linked base was either the 5’ most deletion in the read, or the base immediately 5’ to the read end if no deletion was present. For eIF4A3 and Casc3 iCLIP datasets, obtained reads were already filtered, trimmed and demultiplexed and entered the above process at the mapping stage. We treated each replicate separately, and also report results from a ‘union’ set generated by merging iCLIP tags from all replicates for a factor. All iCLIP track visualizations were prepared using the Integrative Genomic Viewer (Robinson et al., 2011).

#### Identification of significantly crosslinked bases

As our analyses suggested that components of the TREX complex bind in a broad manner and not to specific “binding sites”, most analyses are conducted using all iCLIP reads. Where specifically noted, significant bases were identified using the procedure outlined in (Wang et al., 2010) and implemented in the ‘significant_bases_by_randomisation’ script from iCLIPlib. For each gene we merged all overlapping exons and then divided the gene into a single exonic region and a separate region for each intron. Within each region the height of an individual base is the number of all crosslink bases within 15 nts. For each region we calculate the empirical distribution of base heights, such that *P_h_* is the fraction of bases in a region with *height* > *h*. To calculate an FDR, we randomise the location of crosslinks within the region and calculate *P_h_* for the randomised profile. This procedure is repeated 100 times, and we then calculate the FDR for a base with height ≥ *h* as *FDR*(*h*) = (*µ_h_* + *σ_h_*)/*P_h_* where *µ_h_* and *σ_h_* are the mean and standard deviation of *P_h_* from the randomisations. We selected bases with FDR < 0.1 as significant. Clusters of significant bases were determined by merging significant bases within 15 nts of each other.

#### Searching for enriched kmers

Enrichment for kmer motifs was conducted using the z-score approach as used in (Wang et al., 2010) as implemented by ‘iCLIP_kmer_enrichment’ from iCLIPlib. All exons for each gene were merged. We then calculated the height of each base in a gene as the number of crosslinks within 15nt of the position. For each possible sequence s of length k, we then calculated its frequency as *f_s_* = Σ*_i_*_∈_*_P_ h_i_* where *P* is the set of locations at which a match to the kmer begins (across the whole transcriptome) and *h_i_* is the height at location *i*. We then calculate *z_s_* = (*f_s_* − *µ_s_*)/*σ_s_* where *µ_s_* and *σ_s_* are the mean and standard deviation of *f_s_* from 100 randomisations of the crosslink positions, where randomisations take place within genes. We calculated z-score for k=6 and 7 for each replicate of each factor and for the union of all replicates, using both all iCLIP tags and only significantly crosslinked bases, for both all exonic sequence and for 3’ UTRs only. We noted that z-scores between pulldowns were correlated with z-scores from the controls and both were correlated with the uracil content of the kmer. To account for possible crosslinking preferences, we calculated the distance of the z-score for the kmer in the test protein from a linear regression line of z scores from the test protein against z-score from the control iCLIP.

#### Metagene analysis

Metagene profiles of iCLIP data across genes were obtained using ‘iCLIP_bam2geneprofile’ from iCLIPlib. Expressed (see above) protein coding transcripts were divided into a fixed number of bins and iCLIP tags in each bin summed. Flanking regions were scaled so that bins in the flanks were the same size as the exonic bins. Profiles over each gene were normalised to the sum across all regions (usually upstream, exons, downstream) for that gene. Genes with zero tags were excluded. The final profile was then calculated by summing across all genes. For profiles with separate regions for UTRs and CDS, each region from each expressed protein coding transcript was divided into a number of bins (20, 100 and 70 for 5’ UTR, CDS and 3’ UTR respectively). Tag counts in each bin were calculated and normalised to the size of the bin for that region and that transcript. Counts for each transcript were then normalised to the sum of all normalised bin counts for that transcript. We repeated this process for total nuclear RNA-seq and divided each iCLIP profile by the RNA-seq profile. This controls for miss-annotation of the start of 5’ UTRs and the end of 3’ UTRs. For exon-exon junction profiles, we first excluded the first and last junction in each transcript and then excluded junctions where the exons on either side of the junction were less than 100 nts in length. For each of the remaining junctions we counted the number of iCLIP tags at each position within 100 nts of the junction (in transcript coordinates) and normalised to the total number of iCLIP tags within 100 nts either side of the junction. Profiles were then calculated by summing over all junctions and normalized to the sum of the profile.

#### Enrichments over control

We counted iCLIP tags in each gene and each ‘transcript chunk’ (see above) using ‘count_clip_sites’ from iCLIPlib. The gene biotype was that assigned by Ensembl (see https://www.gencodegenes.org/ for definition of each biotype). Enrichments over control and confidence intervals were calculated using the ‘boot_ci’ function from iCLIPlib. For gene biotypes, enrichment was calculated by summing tag counts for genes in a category (e.g. protein coding genes) and dividing by the sum of tag counts for the control. In order to measure our uncertainty, we took 1000 bootstrap samples of genes and for each bootstrap calculated the ratio of pulldown tag counts to control tag counts. Confidence intervals encompass 95% of the values from these samples. The same process was used for the calculation of enrichment over control for other categories, with the exception that bootstrap samples of exons were taken where categories were categories of exons rather than transcripts (e.g. first, middle, last exons or short vs long exons). For nuclear/cytoplasmic lncRNAs, we took transcripts assigned the lincRNA biotype. To assign transcripts to nuclear or cytoplasmic categories, we generated read counts from the nuclear and cytoplasmic total RNA seq (see above). We assigned a gene to the nuclear category if the expression was significantly higher in the nucleus as measured by DESeq2 (Love et al., 2014) with a 5% FDR cutoff. Enrichments over control and confidence interval was calculated as above for biotypes.

#### Splicing Index

To calculate splicing index, we first collected reads that overlapped introns. We then put reads into one of 4 categories: Spliced (S) reads were those that overlapped both exons flanking the intron and contained a splice at exactly the annotated coordinates; Exon-Intron (EI) reads had at least 3 nts aligned to the 3’ end of the 5’ exon and at least 3 nts aligned to the 5’ end of the intron; Intron-Exon (IE) reads had at least 3 nts aligned to the 3’ end of the intron and at least 3 nts aligned to the 5’ end of the 3’ exon; and “other” reads. The splicing index (SI) was calculated as:

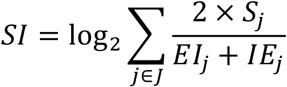

where J is the set of all annotated junctions and *Sj, EI_j_* and *IE_j_* is the count of Spliced, Intron-Exon and Exon-Intron reads at junction j respectively.

#### Processing Index

To calculate the processing index, we obtained HEK293 A-seq data (Martin et al., 2012) and used it to select the 3’ most poly-A site overlapping the expressed transcripts from each gene. For each gene g in the set, we counted the number of iCLIP tags in a 50 nts window upstream (*u_g_*) and downstream (*d_g_*) of each poly-A site. As the signal upstream of the poly-A site originates from both processed and unprocessed transcripts, while that downstream originates only from processed, we estimate counts from processed transcripts as *u_g_* − *d_g_*. Thus, the processing index is:

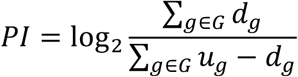

#### Analysis of RNAseq data

Reads for mRNA-seq of siControl and siChtop transfected cells were adaptor and quality trimmed, removing bases with a Q<15 from the 5’ end of the read, Q<20 from the 3’ end of the read and also if the average quality in a 4 nts window falls below 15. Trimmed reads greater than 36 nts were retained.

These reads and ribosome depleted nuclear and cytoplasmic total RNA from PRJEB4197 were mapped against the hg19 genome sequence using STAR 2.4.2a and a junction database built from Ensembl 75. The options to STAR were ‘--outFilterType BySJout --outFilterMismatchNmax 5’. All RNA-seq track visualizations were prepared using the Integrative Genomic Viewer.

#### Alternate splicing of detained and retained introns

We defined two categories of potentially inefficiently spliced introns: Retained and Detained. “Retained introns” are isoforms of a gene where an intron is retained compared to another isoform. Detained introns are as defined by (Boutz et al., 2015) and are introns with higher than usual RNA-seq signal in a region not defined as an exon in any isoform of a gene.

#### Identification of retained introns

To identify retained introns, we took all the isoforms of a gene and compared them pairwise. Where all introns in transcript A are present in transcript B, but transcript B contains introns not present in transcript A that are contained between the start and end of transcript A, we call these “retained introns”.

#### Identification of detained introns

To identify “detained” introns, we applied the method of (Boutz et al., 2015). We compared intronic read counts from nuclear mRNA-seq to counts from *in silico* constructed null repeats. We first counted reads in each transcript chunk that overlaps no exons (i.e. constitutive introns), requiring a 10 nts overlap and disallowing multimapping reads. We then used DESeq2 to calculate library size normalised counts for each of the three replicates. We then used the normalised total of intronic reads from each gene to generate one null replicate for each of the real replicates. For each intron, we calculated a weight based on its effective length:

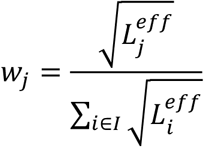

Where *I* is the set of introns in a gene and 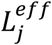 is the effective length of intron j and is equivalent to the number of positions which a valid read alignment could begin from and still be counted as coming from that intron. It is calculated as:

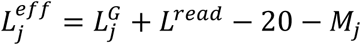

Where 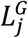 is the genomic length of the intron, 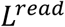 is read length, 20 is twice the amount of overlap required for a read to count and *M_j_* is the number of positions in the intron that would lead to a multimapping read as determined from the 100mer alignability track downloaded from http://genome.ucsc.edu/cgi-bin/hgFileUi?db=hg19&g=wgEncodeMapability. Introns with a weight of 0 were excluded from further consideration. For each replicate, we now distributed the sum of normalized intronic reads between the introns of a gene using the weights. If *C_i,r_* is the normalised counts of reads in intron i in replicate r, then the null counts are:

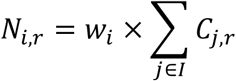

We repeated this process for all three replicates of RNA-seq data and then used DESeq2 to identify introns with a significantly higher than expected read count using an FDR threshold of 1% and only taking those with at least 4 times the expected count.

#### Identification of introns with increased splicing efficiency on Alyref knockdown

To identify introns with increased splicing efficiency using nuclear mRNA-seq of Control and Alyref knockdown cells, we first counted reads that mapped to each ‘chunk’ of a gene (see above) using ‘featureCounts’; counting only those reads that overlapped chunks by at least 10 nts, and keeping multi-mapped reads (but only the primary alignment), that is with the options ‘-O --primary -M --minOverlap 10 -f’. We then applied DEXSeq (Anders et al., 2012) to these counts to identify differential exon (or intron) usage, selecting chunks using a 10% FDR threshold and at least a 1.5 fold increase compared to the rest of the gene. To calculate the Alyref density on each transcript chunk, we took the log_2_ of the number of Alyref iCLIP tags from all replicates divided by the genomic size of the chunk. Chunks of less than 10 bp were excluded from further analysis. We then divided the range of Alyref binding densities into three equally sized bins (Low, Medium and High). We compared the differentially spliced regions to our annotations of retained and detained introns, described above, and calculated the fraction of retained, detained and other introns whose levels had increased compared to other regions of the gene.

##### Analysis of alternative polyadenylation changes

To determine genes with a change in poly-A site usage upon Chtop knockdown, we converted mRNA-seq reads from siControl and siCHTOP transfected HEK293T cells into read depth and used the DaPars program (Xia et al., 2014) to select genes with an absolute change in Percentage of Distal pA site Usage Index (PDUI) of greater than 0.25, with an FDR threshold of 5%. To calculate change in weighted transcript length, we quantified transcript expression using Salmon v0.8.2. We then used the TxImport package (Soneson et al., 2015) to calculate the expression weighted transcript length for each gene. Change in length for each gene was calculated as the log_2_ ratio of mean length in siChtop transfected cells divided by mean length in siControl transfected cells.

### DATA AND SOFTWARE AVAILABILITY

Raw and processed sequencing data have been deposited in the GEO archive with accession GSE113896.

The iCLIPlib software is available from www.github.com/sudlab/iCLIPlib. The precise version used in this manuscript is tagged “Viphakone_et_al_release”. This includes pipeline_iCLIP used for basic iCLIP processing. Code used for further analysis and generation of figures is available at www.github.com/sudlab/Viphakone_et_al.

## SUPPLEMENTAL INFORMATION

Supplemental Information includes six figures and one table.

## ACKNOWLEDGEMENTS

SW acknowledges support from Biotechnology and Biological Sciences Research Council (UK) (grants BB/N014839/1 and BB/N005430/1) and funding from the Medical Research Council via the Computational Genomics analysis and Training Programme (CGAT). We thank Dr. Nicholas Conrad for providing us his mRNA-seq datasets from Control and Alyref knockdown cells. We thank the Wilson Lab members and Dr. Julie Aspen for useful discussions.

## AUTHOR’S CONTRIBUTIONS

N.V. performed all molecular biology, iCLIP, and transcriptomic experiments with help from C.G.H. I.S. designed and performed all bioinformatics analyses with input from N.V., S.A.W, and D.S. N.V. and S.A.W designed the project. N.V. and S.A.W wrote the article with input from all authors.

